# Rac-dependent signaling from keratinocytes promotes differentiation of intradermal white adipocytes

**DOI:** 10.1101/474056

**Authors:** Takehiko Ueyama, Megumi Sakuma, Mio Nakatsuji, Tatsuya Uebi, Takeshi Hamada, Atsu Aiba, Naoaki Saito

**Author notes:** Correspondence and requests for materials should be addressed to TUeyama or NS.

## Abstract

Rac signaling affects numerous downstream targets; however, few studies have established in vivo levels. We generated mice with a single knockout (KO) of *Rac1* (*Keratin5 (K5)-Cre;Rac1^flox/flox^*, *Rac1*-KO) and double KO of *Rac1* and *Rac3* (*K5-Cre;Rac1^flox/flox^;Rac3^−/−^*, *Rac1/Rac3*-DKO) in keratinocytes. Strikingly, *Rac1-KO* mice exhibited thinner dermal white adipose tissue, which was considerably further reduced in *Rac1/Rac3*-DKO mice. DNA microarray using primary keratinocytes from *Rac1/Rac3*-DKO mice exhibited decreased mRNA levels of *Bmp2*, *Bmp5*, *Fgf20*, *Fgf21*, *Fgfbp1*, and *Pdgfα*. Combinational treatment with BMP2 and FGF21 or BMP2 and FGF20 in culture medium, but not individual purified recombinant proteins, could differentiate 3T3-L1 fibroblasts into adipocytes, as could culture media obtained from primary keratinocytes. Conversely, addition of anti-BMP2 or anti-FGF21 antibodies into the culture medium inhibited fibroblast differentiation. Furthermore, combinational treatment with BMP2 and FGF21 promoted adipocyte differentiation only of rat primary white, but not brown, adipocyte precursors. Notably, brown adipogenesis by FGF21 was inhibited by BMP2. Thus, we proposed novel paracrine pathways from keratinocytes to intradermal pre-adipocytes, which function as Rac-dependent modulators of white adipogenesis, but also brown adipogenesis.

## Introduction

The skin functions as a barrier between the organism and the environment (Proksch, Brandner et al., 2008), preventing attack by environmental pathogens, chemicals, and UV along with mechanical injury. It also prevents loss of water and solutes from the inside of the organism. The skin is composed of three layers: the epidermis, dermis, and subcutaneous tissue (Koster & Roop, 2007). Keratinocytes comprise the major cell types of the epidermis, constituting approximately 90% of the epidermal cells (Nestle, Di Meglio et al., 2009). They proliferate in the basal layer and subsequently form stratification consisting of three types of postmitotic cells, spinous, granular, and cornified layers (Koster & Roop, 2007). The physiological barrier of the epidermis is established during embryogenesis and mainly localized in the cornified layer, but also in the lower layers (Koster & Roop, 2007, Proksch et al., 2008). The upper part of the dermis is divided into the papillary dermis and reticular dermis, and the lower part of the dermis contains white adipose tissue (WAT) (Driskell, Lichtenberger et al., 2013).

Recently, it has been proposed that two distinct (developmentally, morphologically, and physiologically) types of WAT exist in the skin: intradermal adipocytes (dermal WAT) beneath the reticular dermis and subcutaneous WAT, which are clearly separated by a striated muscle called the panniculus carnosus in rodents (Driskell et al., 2013). Immature adipocytes in the dermis reportedly promote hair follicle cycling through PDGFα secretion (Festa, Fretz et al., 2011), demonstrating the paracrine signaling from intradermal adipocytes to the epidermis to promote differentiation of the stem cells in the follicular bulge. In addition, the possibility of the reciprocal signaling from hair follicles to the intradermal pre-adipocytes; e.g., crosstalk between cells in the bulge (keratinocytes) and intradermal adipocytes, is suspected (Donati, Proserpio et al., 2014, Jahoda & Christiano, 2011); however, the detailed mechanism has not been elucidated. Furthermore, adipose tissues are classified into both WAT and brown adipose tissue (BAT), with beige/brite (beige) adipocytes, comprising brown adipocytes induced in the WAT, being proposed as a third adipocyte type (Sanchez-Gurmaches, Hung et al., 2016). In particular, brown and beige adipocytes express Ucp1 as a marker (Wang & Seale, 2016). However, the induction mechanisms of white, brown, and beige adipocytes are not yet fully understood.

Racs (Rac1, Rac2, and Rac3) comprise the best characterized members of the Rho family of small guanosine triphosphatases (GTPases), which play fundamental roles in a wide variety of cellular processes including transcriptional regulation, establishment and maintenance of cell polarity, and turnover of actin-based structures (Bosco, Mulloy et al., 2009, Namba, Funahashi et al., 2015). Previously, the hairless phenotype was reported in mice with deletion of Rac1 (Benitah, Frye et al., 2005, Castilho, Squarize et al., 2007, Chrostek, Wu et al., 2006), in which either the keratin 5 (K5) or K14 promoter was used to effect Rac1 deletion in keratinocytes. Benitah et al. (Benitah et al., 2005) reported that loss of Rac1 in keratinocytes prevents the renewal of epidermal stem cells and cause a robust commitment to stem cell differentiation, finally resulting in epidermal failure. In addition, the latter two groups reported the critical role of Rac1 in hair follicle development and maintenance. However, the functions of Rac3 in keratinocytes, and whether Rac3 functions synergistically (Nakamura, Ueyama et al., 2017, Vaghi, Pennucci et al., 2014) or antagonistically (Hajdo-Milasinovic, Ellenbroek et al., 2007) with Rac1 in keratinocytes, remain unknown.

To address these issues, in the present study we generated mice with a single KO of *Rac1* under control of the K5 promoter (*K5-Cre;Rac1^flox/flox^*, hereafter *Rac1*-KO) and double KO of *Rac1* and *Rac3* in keratinocytes (*K5-Cre;Rac1^flox/flox^;Rac3^−/−^*, hereafter *Rac1/Rac3*-DKO). We found that additional deletion of *Rac3* exacerbated the phenotypes observed in *Rac1*-KO mice. Furthermore, we observed thinner dermal WAT in *Rac1*-KO mice, with this effect being markedly exacerbated in *Rac1/Rac3*-DKO mice. Notably, *Bmp2* and *Fgf21* mRNAs levels were decreased in primary *Rac1/Rac3*-DKO keratinocytes. Combinational treatment of BMP2 and FGF21 promoted differentiation of pre-adipocytes into white adipocytes but not brown adipocytes. Thus, we embodied a differentiating signaling phenomenon from keratinocytes to intradermal adipocytes using keratinocyte-specific *Rac1/Rac3*-DKO mice, demonstrating the existence of reciprocal signal/crosstalk between epidermal keratinocytes and intradermal white adipocytes.

## Results

### Expression of *Rac1* and *Rac3* in primary keratinocytes and generation of keratinocyte-specific *Rac1/Rac3*-DKO mice

To examine the Rac isoform expressed in keratinocytes, we performed RT-PCR using primary keratinocytes obtained from wild-type (WT) mice. *Rac1* was the predominant Rac isoform expressed in mouse primary keratinocytes (Figure 1A), although *Rac3* was also expressed at a lesser amount (Figure 1B). Next, the pattern of *K5–Cre*-driven recombination was assessed by X-gal staining using *K5–Cre^+/−^;LacZ^+/−^* mice: progeny of *K5–Cre* mice (Tarutani, Itami et al., 1997) and *CAG–STOP^flox^–LacZ* (Sakai & Miyazaki, 1997) reporter mice. Consistent with previous reports (Tarutani et al., 1997), X-gal positive cells comprised keratinocytes in the interfollicular epidermis, hair follicles, and sebaceous glands of the dorsal skin (Figure 1C). Based on these results, we generated mice with a single knockout (KO) of *Rac1* and double KO (DKO) of *Rac1* and *Rac3* in keratinocytes by crossing *Rac1^flox/flox^* mice (Ishii, Ueyama et al., 2017) or *Rac1^flox/flox^;Rac3^−/−^* mice(Nakamura et al., 2017) with K5*–*Cre mice (Tarutani et al., 1997): hereafter referred to as *K5– Cre;Rac1^flox/flox^* (*Rac1*-KO) mice and *K5–Cre;Rac1^flox/flox^;Rac3^−/−^* (*Rac1/Rac3*-DKO) mice (Figure 1D).

**Figure 1:**
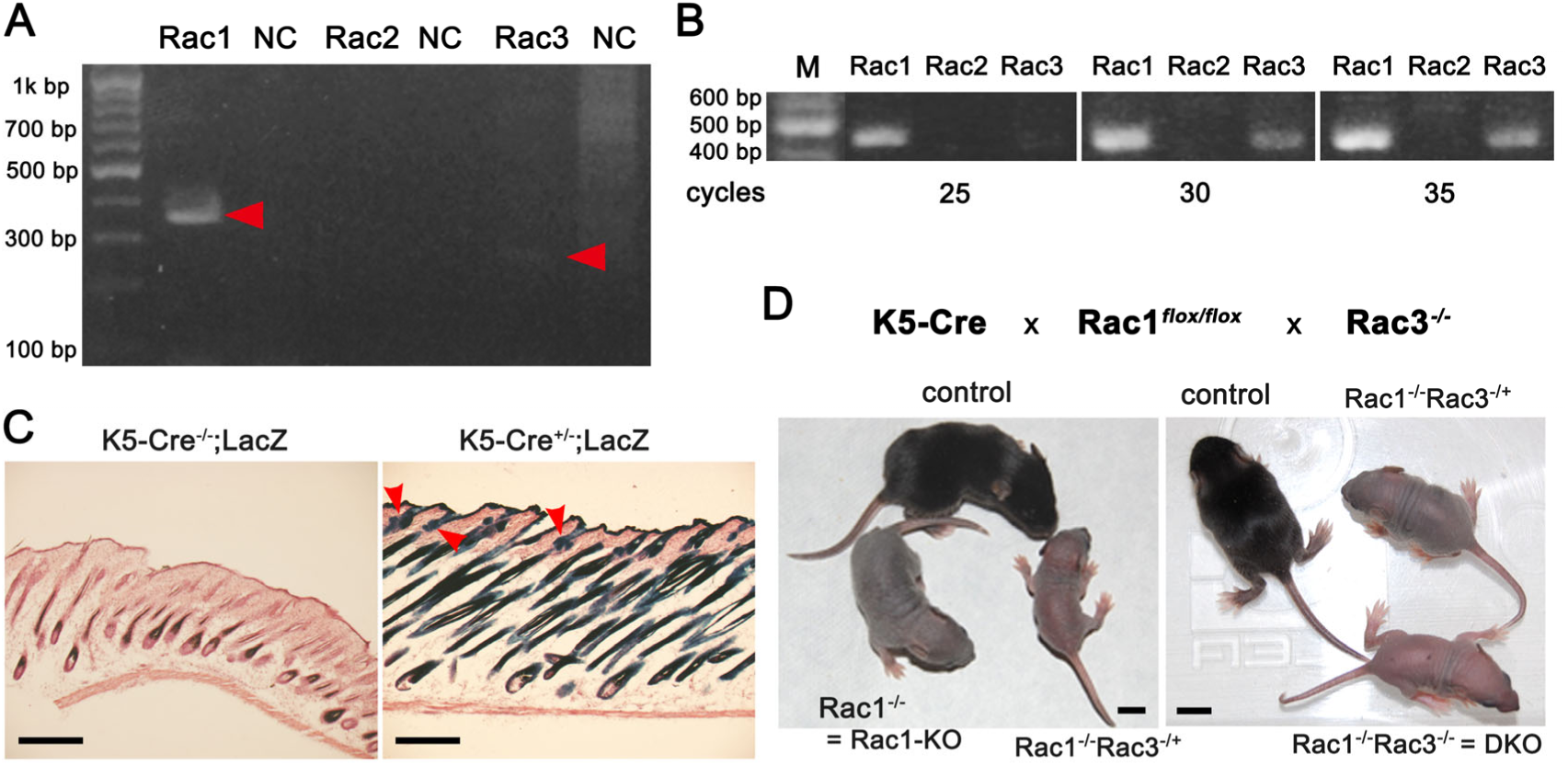
Expression of *Rac1* and *Rac3* in primary keratinocytes and exacerbated phenotypes in Rac1/Rac3-DKO mice. **A** and **B,** RT was performed using mRNAs obtained from P1 WT primary keratinocytes. Subsequently, PCR was performed using specific primer sets for *Rac1*, *Rac2*, and *Rac3* with 30 cycles. In addition to the amplified band of *Rac1* (358 bp), a faint *Rac3* band (257 bp) is observed (**A**). PCR was performed using another specific primer sets for *Rac1*, *Rac2*, and *Rac3* with 25, 30, or 35 cycles. *Rac1* (455 bp) and *Rac3* (441 bp) bands are observed from 25 and 30 cycles, respectively (**B**). **C,** X-gal staining using *K5-Cre^−/−^;LacZ^+/−^* and *K5-Cre^+/−^;LacZ^+/−^* mice at P25. X-gal is positive in the interfollicular epidermis, hair follicle, and sebaceous gland (arrows) of *K5-Cre^+/−^;LacZ^+/−^* mice. Scale bars: 300 μm. **D,** Photographs of control, *K5-Cre;Rac1^flox/flox^* (*Rac1^−/−^*, *Rac1*-KO), *Rac1*-KO with heterozygous *Rac3*-KO mice (*Rac1^−/−^Rac3^−/+^*) (P12, left) and control, *Rac1^−/−^Rac3^−/+^*, K5-Cre;*Rac1^flox/flox^;Rac3^−/−^*(*Rac1^−/−^Rac3^−/−^*, *Rac1/Rac3* double KO (DKO)) mice (P11, right). The strongest and exacerbated hairless phenotype is observed in *Rac1/Rac3*-DKO mice. Scale bars: 10 mm.

### *Rac1/Rac3*-DKO mice exhibit thinner dermal WAT

*Rac1*-KO mice exhibited a hairless phenotype (Benitah et al., 2005, Castilho et al., 2007, Chrostek et al., 2006) whereas *Rac3*-KO mice (*Rac3^−/−^* and *Rac1^flox/flox^;Rac3^−/−^*) showed no phenotype (Cho, Zhang et al., 2005, Corbetta, Gualdoni et al., 2005) (Figure 1D), consistent with previous reports. *Rac3*-KO mice were subsequently used as controls, in addition to *Rac1^flox/flox^* mice. Around postnatal day 10 (P11–12), mice with heterozygous deletion of *Rac3* in additional to Rac1-KO (*K5–Cre;Rac1^flox/flox^;Rac3^−/+^*) showed smaller body size compared with control (*Rac1^flox/flox^*) and *Rac1*-KO mice, and an exacerbated hairless phenotype compared with *Rac1*-KO mice: i.e., *K5– Cre;Rac1^flox/flox^;Rac3^−/+^* mice showed less black color than *Rac1*-KO mice (Figure 1D). Furthermore, *Rac1/Rac3*-DKO mice presented an exacerbated hairless phenotype compared to that of *K5–Cre;Rac1^flox/flox^;Rac3^−/+^* mice, with the former showing relatively less black color (Figure 1D). Although all control (*Rac1^flox/flox^*, *Rac3^−/−^*, *Rac1^flox/flox^;Rac3^−/−^*) mice and all except one *Rac1*-KO mice were alive at P30, *K5–Cre;Rac1^flox/flox^;Rac3^−/+^* mice showed a lower survival rate. Furthermore, all *Rac1/Rac3*-DKO mice died prior to P22, when is before weaning (Figure 2A). Measurement of the body weight of *Rac3*-KO and *Rac1/Rac3*-DKO mice from P2 to P21 revealed that *Rac1/Rac3*-DKO mice exhibited significantly decreased body weight compared with *Rac3*-KO mice from P6 (Figure 2B; WT and *Rac1*-KO mice are shown for reference).

**Figure 2:**
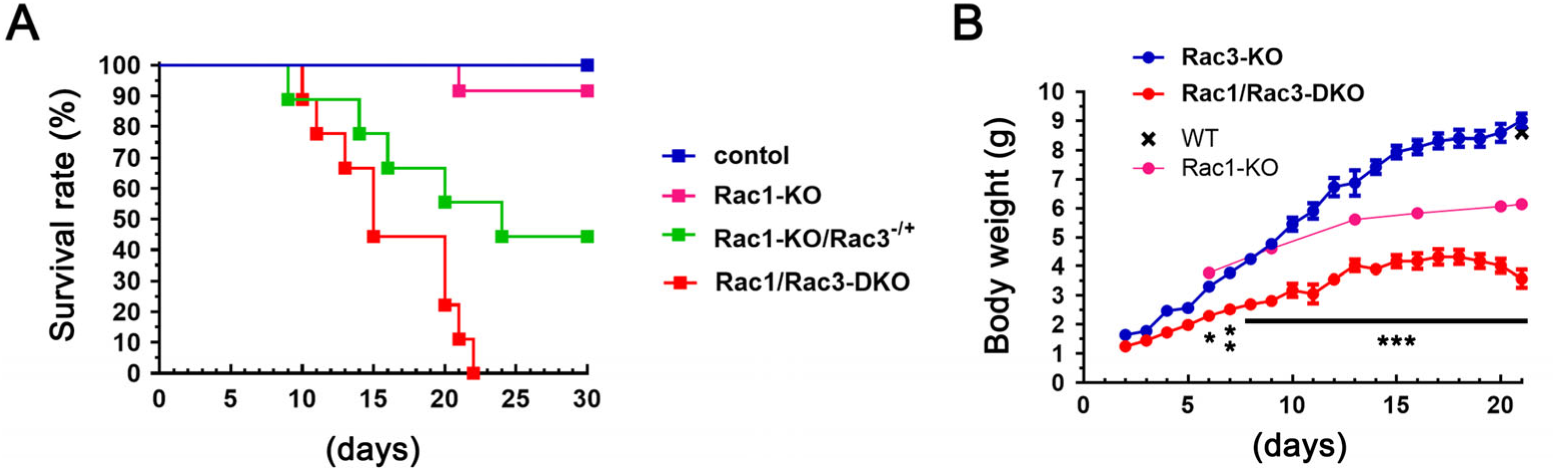
Death and body weight of *Rac1/Rac3*-DKO mice. **A,** Survival rate of control (n = 42: *Rac1^flox/flox^*, *Rac3*-KO, *Rac1^flox/flox^;Rac3^−/−^= Rac3*-KO), *Rac1*-KO (n = 12), *Rac1^−/−^Rac3^−/+^* (n = 9), *Rac1/Rac3*-DKO (n = 9) mice. Only one Rac1-KO mice died prior to P30, and all *Rac1/Rac3*-DKO mice died before P22. **B,** Body weight of *Rac1/Rac3*-DKO mice (n = 11) compared with control mice (*Rac3*-KO, n = 23). Significant differences are observed from P6. \**P*< 0.05, \*\**P* < 0.01, \*\*\**P* < 0.001 by two-way ANOVA followed by Tukey’s *posthoc* test. Rac1-KO (n = 1) and WT (n = 1) are shown for reference.

To clarify the cause of early death of *Rac1/Rac3*-DKO mice, we first examined the permeability (barrier function) of the skin using a toluidine blue assay. No difference in toluidine blue permeability was observed between control and *Rac1*-KO mice (Supplementary Figure S1A). In contrast, *Rac1/Rac3*-DKO mice showed greater permeability of toluidine blue compared with that of control (*Rac3-KO* and *Rac3-KO* plus heterozygous *Rac1-KO*) mice (Supplementary Figure 1B, C). However, this impaired permeability in *Rac1/Rac3-DKO* mice was milder than that in a previous reported mouse line showing no lethality (Sokabe & Tominaga, 2010), and therefore did not appear to be the cause of the early death. We then examined cells types exhibiting a functional *K5* promoter across the whole body by X-gal staining using *K5-Cre^+/−^;LacZ^+/−^* mice. X-gal staining was positive in sweat glands in plantar skin, ependymal cells in the brain, epithelial cells in the esophagus and stomach, and tracheal glands, but negative in the heart, spleen, liver, and kidney (Supplementary Figure 2). In addition, the K5 promoter used in the present study was reportedly very weakly functional in the lung in addition to keratinocytes, esophagus, and stomach (Tarutani et al., 1997); however, we observed that expression in the respiratory tract was limited to just the tracheal glands (Supplementary Figure 2). In the brain, expression was limited to the ependymal cells (Supplementary Figure 2). Thus, we were unable to detect an obvious K5-mediated KO location underlying the early death of *Rac1/Rac3-DKO* mice.

To examine the abnormality in *Rac1/Rac3*-DKO skin, we performed histological examination. The dermis of *Rac1/Rac3*-DKO mice was significantly thin compared with that of the control (*Rac1^flox/flox^*) and *Rac3*-KO (*Rac1^flox/flox^;Rac3^−/−^*) mice from P0 (Figure 3A). In addition, the dermis of *Rac1*-KO mice became thinner than that of *Rac3*-KO mice at P3, and significantly thin compared with that of control (*Rac1^flox/flox^*) and *Rac3*-KO mice at P8 (Figure 3A). From P0, a significant difference of dermal thickness was apparent between *Rac1-KO* and *Rac1/Rac3-DKO* mice (Figure 3A).

**Figure 3:**
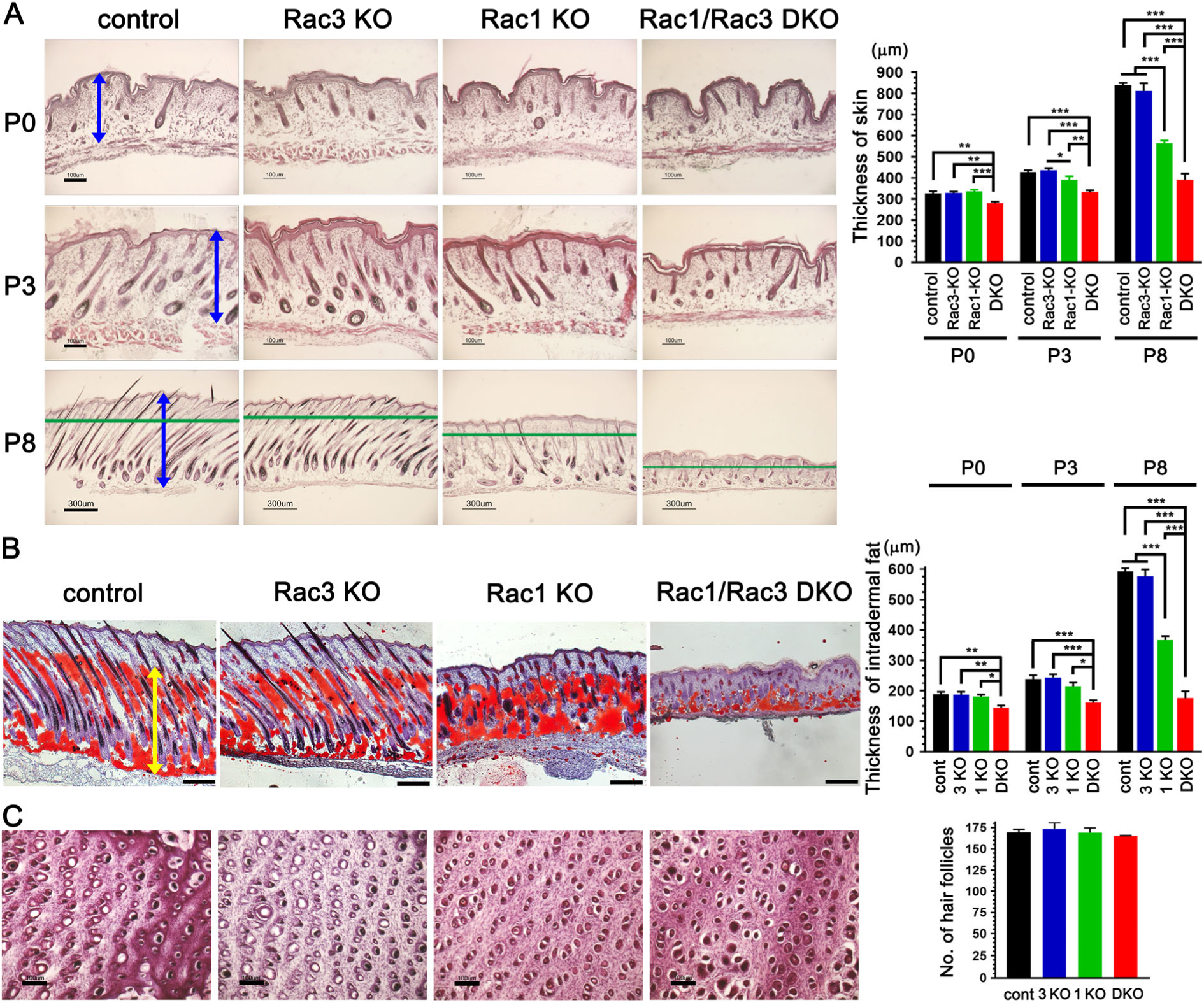
Intradermal fat tissue thickness in Rac1/Rac3-DKO mice. We obtained 20 μm back skin sections from control, *Rac3*-KO, *Rac1*-KO, and *Rac1/Rac3*-DKO mice, and performed HE staining (A and C) and Oil Red O staining (B). **A,** Skin thickness (indicated bidirectional arrows) was measured at P0, P3, and P8, and graphed (n = 4, \**P*< 0.05, \*\**P* < 0.01, \*\*\**P* < 0.001 by one-way ANOVA followed by Tukey’s *posthoc* test). Scale bars: 100 μm (P0 and P3) and 300 μm (P8). **B**, The thickness of Oil Red O positive intradermal fat tissue (indicated bidirectional arrow) was measured at P0, P3, and P8, and graphed (n = 4, \**P*< 0.05, \*\**P* < 0.01, \*\*\**P* < 0.001 by one-way ANOVA followed by Tukey’s *posthoc* test). Oil Red O stained sections obtained from P8 mice are shown. Scale bars: 200 μm. **C,** HE stained perpendicular sections at the level of the infundibula of the hair follicle (indicated by lines in **A** of P8) of control, *Rac3*-KO, *Rac1*-KO, *Rac1/Rac3*-DKO mice at P8 are shown and quantified in a graph (n = 3; no significant difference was observed). Scale bars: 100 μm.

To further examine the most severely affected layer in the dermis, we performed Oil Red O staining, which detects adipocytes. The dermal WAT of *Rac1/Rac3*-DKO mice was significantly thinner than that of control and *Rac3*-KO mice from P0 (Figure 3B). The thickness of the dermal WAT in *Rac1*-KO mice became significantly thinner than that of the control and *Rac3*-KO mice at P8 (Figure 3B). These results are similar to those observed in the skin analysis (Figure 3A), suggesting that the most severely affected layer in the dermis was the dermal WAT. The most notable findings were that the *Rac1/Rac3*-DKO dermal WAT was prevented from increasing its thickness from P0 (no significant difference was observed between P0 vs P3 (*P* = 0.9992), P3 vs P8 (*P* = 0.9998), or P0 vs P8 (*P* = 0.8971) by two-way ANOVA followed by Tukey’s *post hoc* test, Figure 3B). Moreover, although no significant increase was observed between P0 and P3 (*P* = 0.8413) for *Rac1*-KO dermal WAT, a significant increase was observed between P3 and P8 (*P* < 0.001 by two-way ANOVA followed by Tukey’s *post hoc* test, Figure 3B).

The number of the upper portion of hair follicles (infundibula) did not significantly differ among control, *Rac3*-KO, *Rac1*-KO, or *Rac1/Rac3*-DKO mice (Figure 3C). This result was consistent with a report that abnormalities in hair follicles were limited to the nonpermanent part of the hair follicle in *Rac1*-KO mice (Chrostek et al., 2006), which is not associated with the cyclic growth of hair follicles (Fuchs, 2007).

### Reduced secreted factors from *Rac1/Rac3*-DKO keratinocytes

We hypothesized that the thinner dermal WAT in *Rac1/Rac3*-DKO mice is associated with decreased secreted factor(s) from *Rac1/Rac3*-DKO keratinocytes. To identify such secreted factor(s), we performed DNA microarray analysis using mRNAs obtained from primary keratinocytes of control (*Rac3*-KO) and *Rac1/Rac3*-DKO mice (Figure 4A). Among the genes decreased by more than twofold in the *Rac1/Rac3*-DKO keratinocytes relative to the *Rac3*-KO keratinocytes (data are deposited in NCBIs GEO, GES122234), we selected six secreted factors as candidates: Bmp2, Bmp5, Fgf20, Fgf21, Fgfbp1, and Pdgfα (Figure 4A). Reduction of Bmp2, Fgf20, Fgf21, Fgfbp1, and Pdgfα mRNAs was confirmed by RT-PCR using equivalent samples (Figure 4B).

**Figure 4:**
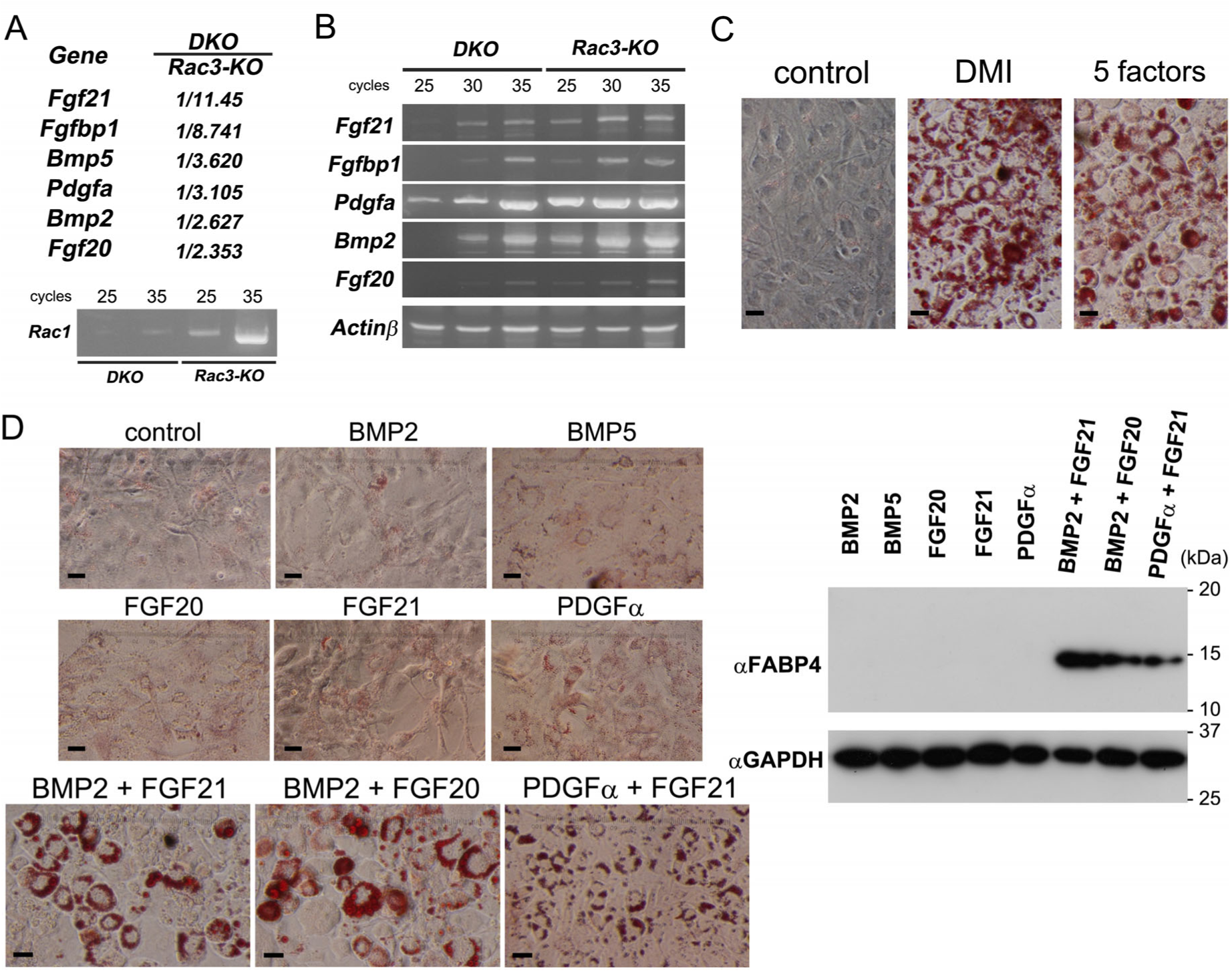
Production of *Bmp2*, *Bmp5*, *Fgf20*, *Fgf21*, and *Pdgfα* from *Rac1/Rac3*-DKO keratinocytes and induction of adipogenesis by BMP2 + FGF21. **A,** DNA microarray analysis was performed using mRNAs obtained from P3 *Rac3* and *Rac1/Rac3*-DKO keratinocytes. *Fgf21*, *Fgfbp1*, *Bmp5*, *Pdgfα*, *Bmp2*, and *Fgf20* were selected as Rac-dependent secreted factors form keratinocytes. Levels of *Rac1* mRNA in *Rac1/Rac3*-DKO and *Rac3*-KO keratinocytes were confirmed by RT-PCR. **B,** mRNA levels of *Fgf21*, *Fgfbp1*, *Pdgfα*, *Bmp2*, and *Fgf20* in Rac1/Rac3-DKO keratinocytes were confirmed by RT-PCR. **C and D,** 3T3-L1 fibroblasts were cultured with treatment of secreted factor(s) for 5 days, followed by Oil Red O staining and immunoblotting. Induction of Oil Red O staining by treatment with DMI or recombinant proteins of the five secreted factors (BMP2, BMP5, FGF20, FGF21, and PDGFα) and control (without treatment) (**C**). Scale bars: 50 μm. Induction of Oil Red O staining by treatment with diluent control, BMP2, BMP5, FGF20, FGF21, or PDGFα, or with BMP2 + FGF21, BMP2 + FGF20, or PDGFα + FGF21 (**D**). 3T3-L1 lysates after treatment with recombinant the secreted factor(s) were subjected to immunoblotting using an anti-FABP4 antibody. FABP4 is observed only in 3T3-L1 cells treated with BMP2 + FGF21, BMP2 + FGF20, or PDGFα + FGF21. Comparable loading proteins were confirmed using an anti-GAPDH antibody.

### Differentiation of 3T3-L1 fibroblasts to adipocytes by BMP2 + FGF21, BMP2 + FGF20, and PDGFα + FGF21

To examine the effects of the six selected factors on adipogenesis, we firstly added purified recombinant proteins of BMP2, BMP5, FGF20, FGF21, and PDGFα (purified recombinant protein of Fgfbp1 was not commercially available) into the culture medium of 3T3-L1 fibroblasts, and examined adipogenesis by Oil Red O staining. 3T3-L1 cells supplemented with the five factors as well as those with dexamethasone, isobutyl-methylxanthine, and insulin (hereafter DMI) as a positive control were positive for Oil Red O staining (Figure 4C). However, none of the single additions of each secreted factor exhibited positive Oil Red O staining (Figure 4D). Next, to define the secreted factor(s) actually involved in 3T3-L1 cell adipogenesis, we analyzed the effect of all combination patterns of two factors (10 patterns) with 3T3-L1 cells. Among these, three patterns of the two-factor combination: BMP2 + FGF21, BMP2 + FGF20, and PDGFα + FGF21, led to positive Oil Red O staining (Figure 4D). Adipogenesis of 3T3-L1 cells mediated by these three combinations was confirmed using immunoblotting for FABP4, a marker of differentiated adipocytes (Figure 4D). Furthermore, adipogenesis induced by BMP2 + FGF21 was also confirmed in C3H/10T1/2 mesenchymal cells (Supplementary Figure 3).

### Induction of adipogenesis of 3T3-L1 fibroblasts by the culture medium from primary keratinocytes

To confirm the effects of secreted factors from keratinocytes on adipogenesis, we used the culture medium of primary keratinocytes. Firstly, we examined the effects of the HuMedia-KG2 based culture medium of mouse primary keratinocytes (MPKs). The culture medium of MPKs cultured for over 48 h (undiluted 48 h cultured MPK medium, 2-fold diluted 72 h cultured MPK medium using fresh Humedia-KG2, and undiluted 72 h cultured MPK medium) promoted the increased expression of FABP4 compared with that of fresh HuMedia-KG2, 24 h cultured MPK medium, 2-fold diluted 48 h cultured MPK medium using fresh HuMedia-KG2, and Dulbecco’s modified Eagle medium (DMEM) containing 10% fetal bovine serum (FBS) (Figure 5A). Next, we examined the effects of HuMedia-KG2 based culture medium of normal human primary epidermal keratinocytes (NHEKs). We decided to focus on FGF21 rather than FGF20, because FGF21 constitutes an endocrine-acting type of FGF, whereas FGF20 represents the paracrine-acting type(Goetz & Mohammadi, 2013). Increased expression of FABP4 protein in 3T3-L1 cells was also observed by the 48 h cultured medium of NHEKs (no dilution), but was inhibited by addition of anti-BMP2 antibodies in the culture medium (Figure 5B). Furthermore, the increased expression of FABP4 protein was decreased by addition of anti-FGF21 antibodies (Figure 5C). These results strongly supported the idea that secreted factors from keratinocytes, especially Bmp2 + Fgf21, promote adipogenesis of the dermal WAT.

**Figure 5:**
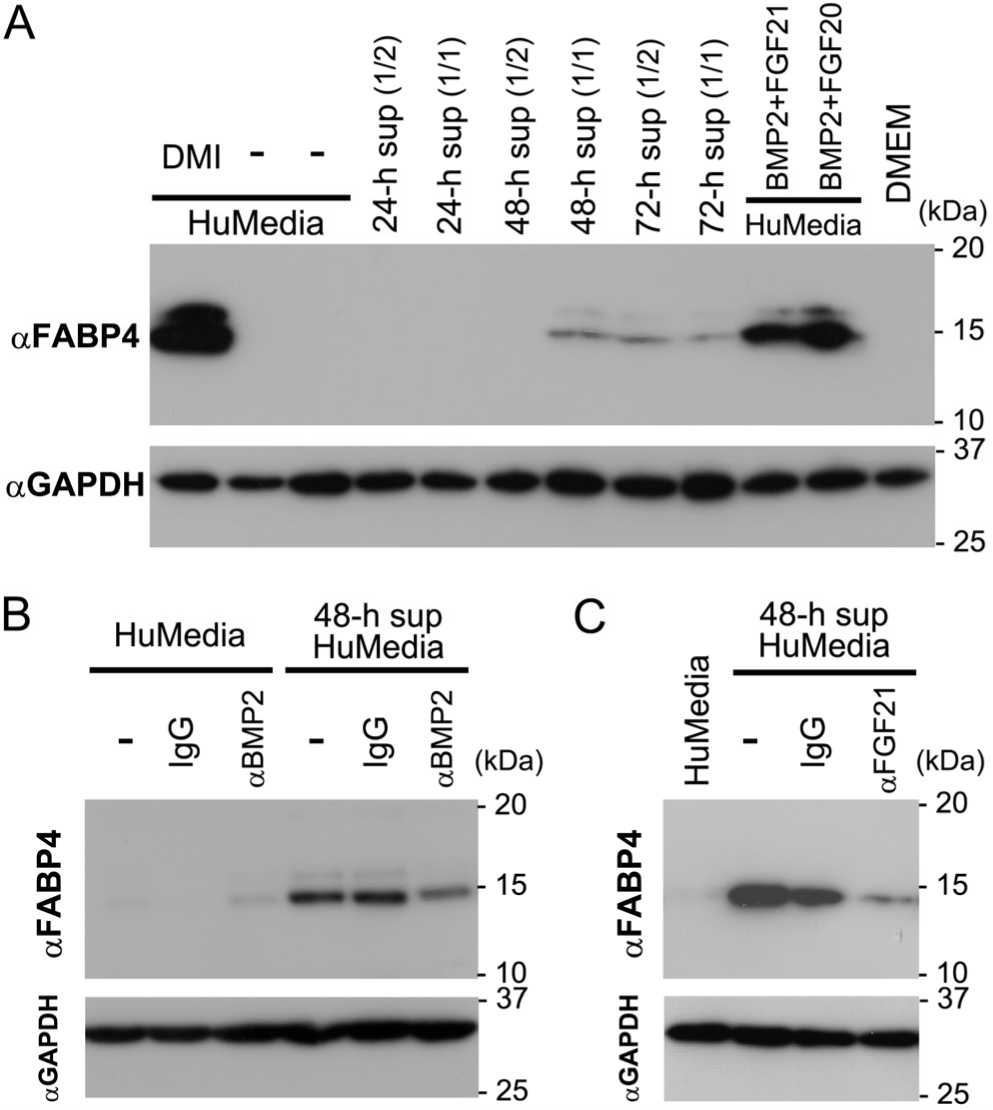
Induction of adipogenesis in 3T3-L1 fibroblasts by culture medium of primary keratinocytes. **A,** 3T3-L1 cells were cultured using Humedia-KG2 (with or without DMI), Humedia-KG2 cultured with mouse primary keratinocytes (MPK) for 24, 48, or 72h (1/2 indicates that cultured Humedia-KG2 with MPK was diluted with the same volume of fresh Humedia-KG2), Humedia-KG2 with BMP2 + FGF21, Humedia-KG2 with BMP2 + FGF20, or DMEM (+FBS). FABP4 immunobands are observed in 3T3-L1 lysates treated with DMI, HuMedia-KG2 cultured with MPK for 48 (1/1) or 72 h (both 1/2 and 1/1), HuMedia-KG2 with BMP2 + FGF21, and HuMedia-KG2 with BMP2 + FGF20. Comparable loading proteins were confirmed using an anti-GAPDH antibody. **B** and **C**, 3T3-L1 cells were cultured using HuMedia-KG2 and Humedia-KG2 cultured with NHEKs for 48 h. FABP4 immunobands induced by Humedia-KG2 cultured with NHEK are reduced by addition of anti-BMP2 antibodies, but not IgG (**B**). FABP4 immunobands are also reduced by addition of anti-FGF21 antibodies, but not IgG (**C**). Comparable loading proteins were confirmed using an anti-GAPDH antibody. sup: supernatant.

### BMP2 + FGF21 induces white, but not brown, adipogenesis

Thin dermal WAT in *Rac1/Rac3*DKO mice and adipocyte differentiation of 3T3-L1 fibroblasts suggested that BMP2 + FGF21 induces Rac-dependent adipogenesis. To further examine which types of adipocytes, i.e., white adipocytes and/or brown adipocytes, are induced by BMP2 + FGF21, we used white adipocyte and brown adipocyte precursor cells, which were isolated from rat abdominal fat tissue and interscapular fat tissue, respectively.

Treatment of white adipocyte precursors with BMP2 + FGF21 induced strong positivity upon Oil Red O staining (Figure 6A): the order of Oil Red O positivity was FGF21 < BMP2 < BMP2 + FGF21. This result was confirmed by immunoblotting of FABP4 (Figure 6B). These results supported that a single treatment of BMP2 may induce white adipogenesis, albeit not to a statistically significant degree; however, combinational treatment of BMP2 + FGF21 markedly enhanced white adipogenesis. Induction of adipogenesis by PDGFα + FGF21 was not observed in white adipocyte precursor cells.

**Figure 6:**
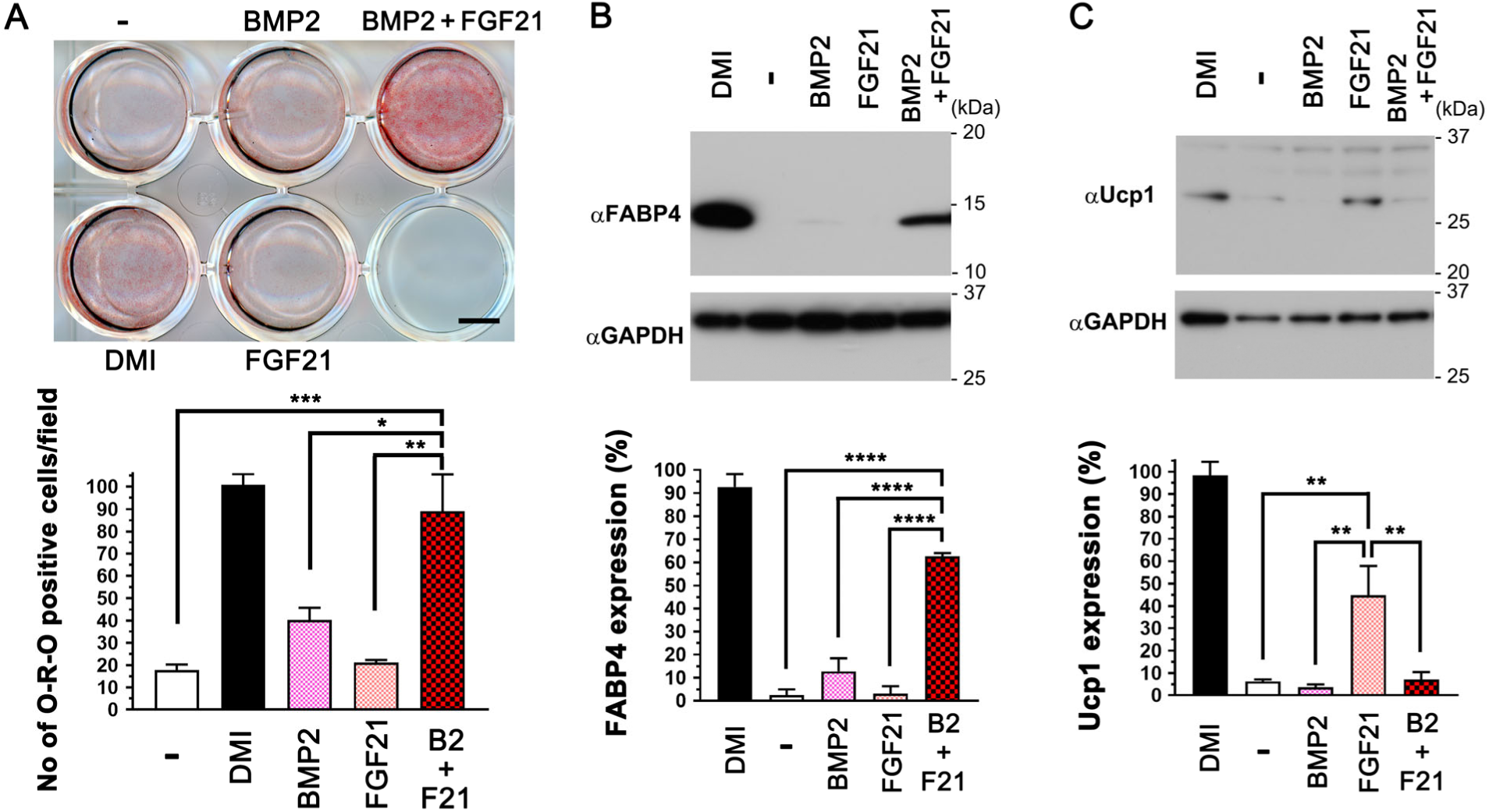
Induction of white or brown adipogenesis by BMP2 + FGF21. White adipocyte precursor (**A, B**) and brown white adipocyte precursor (**C**) cells were cultured with the keratinocyte basic medium supplemented with diluent control (-), DMI, BMP2, FGF21, or BMP2 + FGF21 for ten days. **A,** White adipocyte cells were fixed and subjected to Oil Red O staining. Oil Red O positive cells were quantified and graphed. n = 3, \**P* = 0.0124, \*\**P* = 0.0012, \*\*\**P* = 0.0008 by one-way ANOVA followed by Tukey’s *posthoc* test. Scale bar: 5 mm. **B,** White adipocyte cell lysates were subjected to immunoblotting using an antiFABP4 antibody. FABP4 immunobands were quantified and graphed. Comparable loading of proteins was confirmed using an anti-GAPDH antibody. n = 4, \*\*\*\**P* < 0.0001 by one-way ANOVA followed by Tukey’s *posthoc* test. **C,** Brown adipocyte cell lysates were subjected to immunoblotting using an anti-Ucp1 antibody. Increased levels of Ucp1 are only observed in cells treated with DMI or FGF21, but not BMP2 + FGF21. Comparable loading of proteins was confirmed using an anti-GAPDH antibody. n = 4, \*\**P* < 0.01 (*P* = 0.0074 for control (-) vs. FGF21, *P* = 0.0044 for BMP2 vs. FGF21, and *P* = 0.0088 for BMP2 + FGF21 vs. FGF21) by one-way ANOVA followed by Tukey’s *posthoc* test.

In sharp contrast, treatment of brown adipocyte precursors with BMP2 + FGF21 did not induce increased levels of Ucp1 protein, a marker of brown adipocytes (Figure 6C). Increased levels of Ucp1 protein were only observed in brown adipocyte precursors treated with DMI or FGF21, but not BMP2 or BMP2 + FGF21. Induction of brown adipocytes by FGF21 is consistent with a previous report (Fisher, Kleiner et al., 2012). Notably, induction of Ucp1 protein by FGF21 was inhibited by the addition of BMP2 (BMP2 + FGF21). Taken together, we found that combinational treatment of BMP2 + FGF21 induces white, but not brown, adipogenesis, whereas BMP2 inhibits brown adipogenesis.

## Discussion

In the present study we demonstrated that Rac1 and Rac3 in keratinocytes synergistically function in the development of hair follicles and intradermal adipocytes. To date, three groups have reported keratinocyte-specific *Rac1*-KO mice (Benitah et al., 2005, Castilho et al., 2007, Chrostek et al., 2006), all of which exhibited impaired hair follicle development. In two reports, Rac1 was deleted during embryogenesis using *K5-Cre* (Chrostek et al., 2006) or *K14-Cre* (Castilho et al., 2007) transgenic mice, with the phenotypes observed in *Rac1*-KO mice using *K5-Cre* mice (Chrostek et al., 2006) being consistent with those observed in our *Rac1*-KO mice in the present study. *Rac1*-KO mice generated using *K14-Cre* mice (Castilho et al., 2007) showed more severe phenotypes than those using *K5-Cre* mice (Chrostek et al., 2006), exhibiting lethality prior to P69. Although not confirmed, the authors proposed that the cause of death was related to defective feeding behavior (Castilho et al., 2007). In the present study, although we found that *Rac1/Rac3*-DKO, but not *Rac1*-KO, mice demonstrated mildly impaired barrier function of the epidermis, we consider that this is unlikely to represent the cause of the early death (prior to P22) of *Rac1/Rac3*-DKO mice. In addition, assessment of *K5* promoter function across the whole body by X-gal staining using *K5-Cre^+/−^;LacZ^+/−^* reporter mice also was unable to definitively identify an affected cell type likely to result in the early death of *Rac1/Rac3*-DKO mice, suggesting that the increased lethality may be due to complex factors.

Intrinsic mechanisms including intercellular communication within the epidermis are essential for development and maintenance of the epidermis including hair growth (Fuchs, 2007). In addition, epithelial-mesenchymal interactions are also essential for development of the skin and its appendages, such as hair, mammary glands, and teeth (Hansen, Coggle et al., 1984, Katagiri & Watabe, 2016, Mikkola & Millar, 2006). Thickness of the dermal WAT continuously cycles in accordance with the hair follicle growth cycle, being thinnest in the telogen phase and thickest in the anagen phase (Fuchs, 2007, Hansen et al., 1984, Schneider, Schmidt-Ullrich et al., 2009), which suggests the presence of signaling from hair follicles (keratinocytes) to intradermal adipocytes. The converse, signaling from intradermal adipocytes to hair follicles (epidermal stem cells) have been reported by several groups (Chen, Smith et al., 2002, Festa et al., 2011, Jong, Gijbels et al., 1998). It has been established in mice that the downgrowth and morphogenesis of hair follicles are complete at P8 (Blanpain & Fuchs, 2006) and approximately P17(Muller-Rover, Handjiski et al., 2001), respectively, which have been misinterpreted as “first anagen” (Muller-Rover et al., 2001). Subsequently, the first hair follicle cycling, which begins from catagen, is initiated (Muller-Rover et al., 2001), and the first actual anagen commences at 4 weeks after birth (Schneider et al., 2009). In the keratinocyte-specific *Rac1/Rac3*-DKO mice generated in the present study, the skin failed to execute the final differentiation necessary to enter the hair growth cycle and increase the thickness of the dermal WAT. Thus, our *Rac1/Rac3*-DKO mice embodied the signaling from keratinocytes to intradermal pre-adipocytes, confirming the existence of reciprocal signal/crosstalk between keratinocytes and intradermal adipocytes (Jahoda & Christiano, 2011).

Bone morphogenic proteins (Bmps) belong to the transforming growth factor-β (Tgf-β) family. Over a dozen molecules in vertebrates have been classified into the Bmp subfamily, which have been further classified into several subgroups including the Bmp2/4, Bmp5/6/7/8, Bmp9/10, and Bmp12/13/14 (Katagiri & Watabe, 2016). These four subgroups of Bmps activate Bmp type I receptors along with Smad1, 5, and 8 (receptor-regulated Smads: R-Smad), and the complex of R-Smads and Smad4 (common partner Smads: co-Smad) translocates into the nucleus to regulate target gene transcription (Miyazono, Kamiya et al., 2010). In general, Bmp2 and Bmp4 have been reported to be involved in white adipogenesis, in contrast to the involvement of Bmp7 in brown adipogenesis (Cristancho & Lazar, 2011, Katagiri & Watabe, 2016, Zhang, Schulz et al., 2010). More recently, Bmp4 has also been reported to induce brown adipogenesis and browning of white adipocytes (Elsen, Raschke et al., 2014, Qian, Tang et al., 2013, Xue, Wan et al., 2014). Although Bmp6 was reported to serve as an inducer of white adipogenesis (Donati et al., 2014), Bmp6 was not identified as a downstream target of Rac signaling in the present study. In addition, whereas Bmp2 and Bmp5 were detected as Rac1 downstream targets in keratinocytes in the present study using DNA microarray analysis, Bmp5 did not induce the adipogenesis of 3T3-L1 fibroblasts even in combination with FGF21 or FGF20. However, we confirmed that Bmp signaling was coordinate with reduced phosphorylation levels of Smad1/5 in the intradermal adipocytes of Rac1/Rac3-DKO mice (Supplementary Figure S1C).

Fibroblast growth factors (FGFs) also constitute secreted proteins that act in an intracrine, paracrine or endocrine fashion. The mammalian FGF family comprises 22 members, which are grouped into one intracrine-acting subfamily, five paracrine-acting subfamilies, and one endocrine-acting subfamily (Goetz & Mohammadi, 2013). FGF20 and FGF21 are grouped into one of the paracrine-acting subfamilies, the FGF9 subfamily, consisting of FGF9, FGF16, and FGF20, and the endocrine-acting subfamily, the FGF19 subfamily consisting of FGF19/15, FGF21, and FGF23, respectively. FGF21 regulates the metabolism of glucose, carbohydrates, and lipids (Kurosu, Choi et al., 2007), and induces brown adipogenesis including browning in WAT (Dutchak, Katafuchi et al., 2012, Fisher et al., 2012, Wei, Dutchak et al., 2012), whereas FGF20 is reportedly involved in the development of hair follicles (Huh, Narhi et al., 2013). Periodic expression of FGF21 and FGF 20 in the skin, which depends on the hair growth cycle, has been reported (Kawano, Komi-Kuramochi et al., 2005). In turn, Fgfbps comprise a family of three secreted proteins that act as Fgf chaperons. Fgfbp1 is the best characterized member in Fgfbps, being shown to bind to Fgf1 and 2 (Fgf1 subfamily) and 7, 10, and 22 (Fgf7 subfamily) (Taetzsch, Brayman et al., 2018). As Fgf21 is a member of endocrine-acting Fgfs, it does not appear to require Fgfbps to reach its place of action. However, Fgf20 is a paracrine-acting Fgf; thus, Fgfbps may function in the delivery of Fgf20 from keratinocytes to intradermal adipocytes by providing protection from trapping and proteolysis in the extracellular matrix. In the present study, we detected decreased levels of *Fgfbp1* mRNA in primary keratinocytes; however, the binding capabilities of Fgfbp1 to Fgf20 and Fgf21 have not been determined. Specifically, the lack of available reagents for Fgfbp1, such as antibodies and recombinant proteins, precluded further examination of the role of Fgfbp1 in adipogenesis in the present study.

Notably, a constitutive active mutant of Rac1 and RhoGDIβ, an inhibitor of the Rho-family small GTPases, were reported to promote and prevent BMP4-induced adipogenesis in C3H10T1/2 mesenchymal cells, respectively (Huang, Zhang et al., 2015). In addition, signaling by the canonical Wnt, Wnt/β-catenin, in the epidermis was shown to promote adipogenesis in addition to epidermal development (Blanpain & Fuchs, 2006, Donati et al., 2014), in which BMP2 and BMP6 (Donati et al., 2014), and FGF20 (Huh et al., 2013) were secreted from keratinocytes and hair follicles following activation of Wnt/β-catenin signaling, respectively. Furthermore, involvement of Rac1 in the Wnt/β-catenin signaling has also been reported (Wu, Tu et al., 2008). Taken together, these reports and our present findings of reduced levels of *Bmp2* and *Fgf20*/*Fgf21* mRNAs in *Rac1/Rac3*-KOD keratinocytes support the possibility that Rac involvement in the Wnt/β-catenin signaling (Wnt/β-catenin → Rac) in keratinocytes promotes intradermal adipogenesis through secretion of Bmp2 and Fgf20/Fgf21 from keratinocytes. Alternatively, Bmp2 was also shown to promote osteoblast and chondrocyte differentiation (Katagiri & Watabe, 2016), with low and high concentrations favoring adipogenesis and osteogenesis, respectively (Wang, Israel et al., 1993). Inhibition of Rac1 promoted the Bmp2-induced osteoblastic differentiation, where phosphorylation levels of Smad1/5 (Bmp2-Smad signaling) were not affected (Onishi, Fujita et al., 2013). In addition, inhibition of Rac was conversely reported to induce adipogenesis in 3T3-L1 cells (Liu, DeYoung et al., 2005). These reports suggest that Rac1 activity constitutes an important element for the commitment to adipogenesis and that other factors, such as condition and cell type, may influence the adipogenesis process.

BMPs, which have been reported to be cyclically secreted mainly from intradermal adipocytes underlying the hair follicles, comprise known inhibitors of hair follicles (Plikus, Mayer et al., 2008). However, secretion of Bmp2 and Bmp4 from keratinocytes was also reported (Jamora, DasGupta et al., 2003). In the present study, we found that Bmp2 and Fgf21 secreted from keratinocytes in a Rac-dependent manner induced white adipogenesis. We also demonstrated that Fgf21, which is well-recognized as a brown adipogenesis inducer, facilitates white adipogenesis if applied in conjunction with Bmp2. Furthermore, we identified a novel function of Bmp2, whereby Bmp2 inhibits brown adipogenesis. Thus, baseline secretion of Bmp2 plus endocrine Fgf21 (or paracrine Fgf20) may play important roles for signaling from keratinocytes to intradermal adipocytes to induce white adipogenesis. Our findings are potentially of use for pro-weight gain and anti-obesity applications, through the converse modulation of white and brown adipocytes by Bmp2 induction (increase/decrease) and reduction (decrease/increase), respectively.

## Methods

### Antibodies

The monoclonal antibodies (mAbs) against FABP4(D25B3)XP (1/1000, RRID:AB_2278257), Smad1(D59D7)XP (1/500, RRID:AB_10858882), and phospho-Smad1/5(Ser463/465)(41D10) (1/500, RRID:AB_491015) were purchased from Cell Signaling Technology. The mAbs against human BMP2 (10 μg/ml, 100230, RRID:AB_2065677) and human FGF21 (10 μg/ml, 461804, RRID:AB_2104486) were from R&D Systems. The polyclonal Ab against Ucp1 (1/500, ab10983, RRID:AB_2241462) was from Abcam. The mAbs against GAPDH (1/20000, RRID:AB_10699462) and tubulin-α (1/3000, RRID:AB_10695326), which are conjugated with HRP, were from MBL International (Japan).

### RT-PCR

RT-PCR was performed with 1 µg of total RNA obtained from keratinocytes of control (Rac3) and Rac1/Rac3-DKO mice using SuperScript III reverse transcriptase and random primers (Invitrogen). The following primer pairs were used for PCR: 5′-GCA GAC AGA CGT GTT CTT AAT TTG C-3′ (forward) and 5′-CAA CAG CAG GCA TTT TCT CTT CC-3′ (reverse, 358 bp predicted band) or 5′-TGT AAC AAA AAC TTG GCA TCA AAT GCG-3′ (reverse, 455 bp) for *Rac1*, 5′-GGA GGA CTA TGA CCG CCT C-3′ (forward) and 5′-GCG CTT CTG CTG TCG TGT G-3′ (reverse, 379 bp) or 5′-AAA TAG GAT GTG GCC TAT GAA CAT CC-3′ (reverse, 582 bp) for *Rac2*, 5′-CCC ACA CAC ACC CAT CCT TC-3′ (forward) and 5′-CAG TGC ACT TCT TGC CTG GC-3′ (reverse, 257 bp) or 5′-TGG AGC TAT ATC CCA GAA AAA GGA G-3′ (reverse, 441 bp) for *Rac3*, 5′-CCT GAT GGA ATG GAT GAG ATC TA-3′ and 5′-TCA GGA CGC ATA GCT GGG GCT TCG G-3′ for *Fgf21* (637 bp), 5′-CGT GAT GAG ACT CCA CAG CCT CA-3′ and 5′-TTA GCA TGA TGT CGC CTG TAA CAT G-3′ for *Fgfbp1* (760 bp), 5′-CGC GAT GAG GAC CTG GGC TTG CC-3′ and 5′-TCA CCT CAC ATC TGT CTC CTC CTC CC-3′ for *Pdgf-alpha* (569 bp), 5′-ACC ATG GTG GCC GGG ACC CGC-3′ and 5′-CTA ACG ACA CCC GCA GCC CTC CAC AA-3′ for *Bmp2* (1188 bp), 5′-TCC ATG GCT CCC TTG ACC GAA G-3′ and 5′-TCA AGT GTA CAT CAG TAG GTC TTT G-3′ for *Fgf20* (639 bp), and 5′-TGT TAC CAA CTG GGA CGA CA-3′ and 5′-TTT GAT GTC ACG CAC GAT TT-3′ for beta-actin (415 bp).

### Animals

All animal experiments were conducted in accordance with the guidelines of Kobe University. The *Rac1^flox/flox^* (Ishii et al., 2017) and *Rac1^flox/flox^;Rac3^−/−^* mice (Nakamura et al., 2017), *K5*-Cre transgenic mice (Tarutani et al., 1997), and *CAG-CAT^flox^-LacZ* mice (Sakai & Miyazaki, 1997) have been described previously. The *K5-Cre;Rac1^flox/+^* progeny of *K5-Cre* and *Rac1^flox/flox^* mice were backcrossed with *Rac1^flox/flox^* mice to obtain *K5-Cre;Rac1^flox/flox^* (herein referred to as *Rac1*-KO) mice. The *K5-Cre;Rac1^flox/+^;Rac3^−/+^* progeny of *K5-Cre* and *Rac1^flox/flox^;Rac3^−/−^* mice were backcrossed with *Rac1^flox/flox^;Rac3^−/−^* mice to obtain *K5-Cre;Rac1^flox/flox^;Rac3^−/−^* (herein referred to as *Rac1/Rac3*-DKO) mice. These mice were in turn backcrossed to *Rac1^flox/flox^* or *Rac1^flox/flox^;Rac3^−/−^* mice to generate experimental animals. *CAG-CAT^flox^-LacZ* mice were backcrossed with *K5-Cre* mice to generate *K5-Cre;LacZ* mice, which were then used to examine the effect of the K5 promoter. The offspring of these mice were genotyped by PCR using the following primers: 5′-ACT CCT TCA TAA AGC CCT CG-3′ (forward) and 5′-ATC ACT CGT TGC ATC GAC CG-3′ (reverse) for *K5-Cre* 5′-ATT TTC TAG ATT CCA CTT GTG AAC-3′ and 5′-ATC CCT ACT TCC TTC CAA CTC-3′ for *Rac1^flox^*, 5′-CAT TTC TGT GGC GTC GCC AAC-3′ and 5′-TTG CTG GTG TCC AGA CCA AT-3′ for *Rac3*^−^, and 5′-CAT TTC TGT GGC GTC GCC AAC-3′ and 5′-CAC GCG GCC GAG CTG TGG TG-3′ for *Rac3^+^*. WT C57BL/6 mice were purchased from CLEA Japan. All mice were identified by numbered ear tags. Mice were housed in specific pathogen-free conditions using an individually ventilated cage system (Techniplast, Tokyo, Japan), and allowed food and water *ad libitum*. The animal facility was maintained on a 14 h light and 10 h dark cycle at 23 ± 2 °C and 50 ± 10% humidity. Mice from the control group were always treated and assessed first, followed by the experimental group. Both male and female mice were used in this study.

### Monitoring of body weight

The weight of mice (*Rac3*-KO vs *Rac1/Rac3*-DKO) was monitored every day from P2 until the death of *Rac1/Rac3*-DKO mice.

### Toluidine blue assay of newborn mice

P0 mice were euthanized using carbon dioxide for 20 min. Mice were sequentially dehydrated in 25, 50, 75, and 100% methanol for 1 min each, and then immersed in 0.1% toluidine blue O (Chroma Gesellschaft Schmidt & Co.) in phosphate buffered saline (PBS)(−) for 20 h at 25 °C. After twice washing using PBS(−) for 20 min, mice were photographed.

### Section preparation and X-gal staining

Animals were deeply anesthetized using pentobarbital, then transcardially perfused with ice-cold 0.9% saline solution and subsequently with 4% paraformaldehyde (PFA) in 0.1 M PB (pH 7.4) (Ueyama, Ninoyu et al., 2016). Dorsal skins were dissected and post-fixed overnight in the same fresh fixative, then 20 μm sagittal sections of the skin were obtained. Hematoxylin and eosin (HE, Muto Pure Chemicals, Japan) staining and X-gal staining counterstained with HE has previously been described (Ueyama, Sakaguchi et al., 2014). For Oil Red O staining, dorsal skins were dissected after transcardial perfusion with ice-cold 0.9% saline solution. Sagittal cryosections (20 μm) of the skin were obtained. After fixation with 4% PFA in 0.1 M PB for 15 min at 23 °C, Oil Red O staining and quantification were performed as described below. The slides were photographed using a light microscope (Axioplan II; Carl Zeiss) and a DP26 camera (Olympus, Japan).

### Isolation of mouse primary epidermal keratinocytes (MEKs)

For isolation of MEKs, back skin was removed from P3 pups. The skins were rinsed with Hank’s balanced salt solution without calcium and magnesium (Wako Pure Chemicals, Japan) and placed in 0.05% collagenase (Sigma-Aldrich) overnight at 4 °C. The epidermis was peeled away from the dermis with forceps. Keratinocytes were detached from the epidermis by stirring and from the dermis by pipetting (Tsunenaga, Kohno et al., 1994). After filtration using a 75 μm cell strainer (BD Biosciences), filtrated cells were gathered by centrifugation at 750 × *g* for 5 min and used for primary culture of keratinocytes, total RNA isolation, and immunoblotting.

### DNA microarray

Total RNA of primary keratinocytes was extracted from P3 *Rac3*-KO and *Rac1/Rac3*-DKO mice using NucleoSpin RNA (MACHEREY-NAGEL GmbH & Co. KG). The quality and quantity of RNA were determined using an Agilent 2100 BioAnalyzer, and gene expression profiles were examined using the SurePrint G3 Mouse GE 8 × 60K Microarray kit (Agilent Technologies). The background-subtracted signal intensity was subjected to 75th percentile normalization for inter-array comparison. The value was used as normalized signal intensity and the ratio of the normalized value of each gene (DKD/*Rac3*-KO) is presented.

### Cell cultures

MEKs from P3 and NHEKs from neonatal foreskin (Kurabo Industries, Japan) were plated on collagen-coated 6 well-plates (Corning), and grown in HuMedia-KG2 supplemented with 0.03 mM Ca^2+^ and 0.1 ng/ml human epidermal growth factor, 10 μg/ml insulin, 0.5 μg/ml hydrocortisone, 0.4% bovine brain pituitary extract, 50 μg/ml gentamycin, and 50 ng/ml amphotericin B (hereafter HuMedia-KG2, Kurabo) at 37 °C in 5% CO_2_.

3T3-L1 fibroblasts (RRID:CVCL_0123) and C3H/10T1/2 mesenchymal cells RRID:CVCL_0190) were purchased from Japanese Collection of Research Bioresources (JCRB) Cell Bank (Ibaragi, Japan), and cultured in DMEM (Wako) containing 10% FBS (Nichirei, Japan).

### Differentiation of 3T3-L1 and C3H/10T1/2 cells to adipocytes by secreted factors

3T3-L1 fibroblasts were cultured in 24 well plates (Falcon) using adipocyte basic medium prepared from DMEM (Wako) supplemented with MK425 (TaKaRa, Japan), which contains ascorbic acid, biotin, pantothenic acid, triiodothyronine, octanoic acid, FBS, and penicillin/streptomycin. To induce differentiation, 2 days post-confluent 3T3-L1 fibroblasts were treated with the following reagents by addition to the culture medium for 5 days: 2.5 μM dexamethasone, 500 μM isobutyl-methylxanthine, and 10 μg/ml insulin (herein DMI) as a positive control; 10 ng/ml of five secreted factors [recombinant human BMP2, recombinant human BMP5, recombinant human FGF20, recombinant human FGF21, and recombinant human PDGF-AA (all from Wako)]; 10 ng/ml of all (10) combination patterns of two secreted factors (including BMP2 + FGF21, BMP2 + FGF20, and PDGF-AA + FGF21); or 10 ng/ml of a single secreted factor (total five patterns).

For differentiation of C3H/10T1/2 mesenchymal cells to adipocytes, confluent cells were firstly pre-cultured in the DMEM containing 10% FBS with or without the following secreted factor(s) (50 ng/ml of BMP2, FGF21, or BMP2 + FGF21) for 6 days, secondary in culture media containing DMI with or without the same secreted factor(s) for 3 days, and finally cultured for 6 days with or without the same secreted factor(s).

### Differentiation of 3T3-L1 cells to adipocytes using culture media of primary keratinocytes

Culture media of MEKs and NHEKs (HuMedia-KG2) were collected at 24, 28, and 72 h after incubation with primary keratinocytes, then centrifuged at 5,000 × *g*. Culture medium of 3T3-L1 cells confluent on 12 well plates was exchanged with 700 μl of HuMedia-KG2 with DMI, HuMedia-KG2, the MEK supernatants (24, 48, and 72 h), the MEK supernatants (24, 48, and 72 h) diluted 2-fold using fresh HuMedia-KG2, HuMedia-KG2 with BMP2 + FGF21, HuMedia-KG2 with BMP2 + FGF20, or DMEM with 10% FBS, and incubated for 5 days. In another experiment, confluent 3T3-L1 cells on 12 well plates were cultured for 5 days using the fresh HuMedia-KG2 or supernatants 48 h cultured with NHEKs. mAb against human BMP2 or human FGF21, or control mouse IgG was added to a concentration of 10 μg/ml in the culture medium. Lysates were subjected to sodium dodecyl sulfate polyacrylamide gel electrophoresis (SDS-PAGE) followed by immunoblotting using a FABP4 Ab.

### Differentiation of rat primary white pre-adipocytes and rat primary brown pre-adipocytes to adipocytes using secreted factors

Rat primary white pre-adipocytes and rat primary brown pre-adipocytes were purchased from TaKaRa Bio Inc. and Cosmo Bio Co. (Japan), respectively. Cells were cultured in 24 well plates (Falcon) using adipocyte basic medium prepared from DMEM (Wako) supplemented with MK425 (TaKaRa), according to manufacturer protocol as described above. After reaching 95% confluence, culture medium was changed into the new adipocyte basic culture medium supplemented with no (as a negative control), or 200 μg/ml of BMP2, FGF21, or BMP2 + FGF21, and cultured for 10 days with an additional change of the medium (once per 5 days). For the positive control of adipocyte differentiation, DMI was added in the adipocyte basic medium for 3 days, and then maintained in the adipocyte basic medium for the same period (10 days). Cells were used for Oil Red O staining and immunoblotting.

### Immunoblotting

Cells were lysed in homogenizing buffer (Ueyama, Tatsuno et al., 2007) by sonication in the presence of a protease inhibitor cocktail, a protein phosphatase inhibitor cocktail (Nacalai Tesque, Japan), and 1% Triton X-100. Total lysates were centrifuged at 800 × *g* for 5 min at 4 °C, then the supernatants were subjected to SDS-PAGE, followed by immunoblotting for 16 h at 4 °C using primary Abs diluted in PBS containing 0.03% Triton X-100 (PBS-0.03T). The bound primary Abs were detected using secondary Ab-HRP conjugates using the ECL detection system (Bio-Rad Laboratories). For quantitative assessment of immunoreactive bands, ImageJ software (National Institutes of Health) was used as previously described (Ueyama et al., 2016). Protein expression levels were normalized to those of GAPDH.

### Oil Red O staining

3T3-L1 adipocytes and rat primary adipocytes cultured in plates were washed with PBS(−) and then fixed with 4% PFA in 0.1 M PB for 20 min at 23 °C. After washing with 60% isopropanol, the fixed cells and skin sections were stained with Oil Red O solution (Muto) for 10 min at 23 °C, which was diluted with water and then filtered. Cells and sections were washed three times with water, then photographed using a camera (EOS Kiss X6i, Canon, Japan) equipped with a microscope (CKX41, Olympus) and scanner (GT-980, Epson, Japan). For quantitative analysis, the number of Oil Red O positive cells/field using a ×20 lens (10 fields per sample) were counted in plates and the thickness of intradermal adipose tissue was measured in sections.

### Statistics

All data are presented as the means ± SEM. Two groups were compared using unpaired Student’s *t*-tests. For comparisons of more than two groups, one-way analysis of variance (ANOVA) or two-way ANOVA was performed followed by Tukey’s *post hoc* test of pairwise group differences. Statistical analyses were performed using Prism 6.0 software (GraphPad).

### Data availability

DNA microarray data are available in the NCBI Gene Expression Omnibus (GEO) under accession number GES122234.

## Acknowledgements

We thank to Prof. Ivan de Curtis (San Raffaele Scientific Institute, Italy) for providing the *Rac3*-KO mice. We are also thankful to Profs. Junji Takeda and Jun-ichi Miyazaki (Osaka University, Japan) for providing the *K5-Cre* mice and *CAG-CAT^flox^-LacZ* mice, respectively. This study was supported by grants from the JSPS KAKENHI program, JP17H04042 to TUeyama, JP17H0432 to NS, and JP26670124 to NS and TUeyama; by grants from the Hyogo Science and Technology Association (26087 and to 30075 TUeyama); and by a grant from the Naito Foundation (2018380 to TUeyama).

## Author contributions

TUeyama planned the project and wrote the manuscript. MS performed experiments using 3T3-L1 cells and primary keratinocytes. MN performed histological experiments. TUeyama and TH performed experiments using primary pre-adipocytes. TUebi monitored mouse body weights and performed experiments using C3H/10T1/2 cells. AA provided the animals. TUeyama and NS analyzed the data.

## Expanded View figure legends

**Supplementary Figure 1: Barrier function in *Rac1/Rac3*-DKO mice.**

Toluidine blue assay was performed at P0 using control (*K5-Cre^−/−^;Rac1^flox/flox^* and *K5-Cre^+/−^;Rac1^flox/+^*) and *K5-Cre^+/−^;Rac1^flox/flox^* (*Rac1*-KO) mice (**A**), and control (*K5-Cre^−/−^;Rac1^flox/flox^;Rac3^−/−^*, *K5-Cre^−/−^;Rac1^flox/+^;Rac3^−/−^*, *K5-Cre^+/−^;Rac1^flox/+^;Rac3^−/−^*) and *K5-Cre^+/−^;Rac1^flox/flox^;Rac3^−/−^* (*Rac1/Rac3*-DKO) mice (**B**). Scale bars: 10 mm (**A** and **B**) and 500 μm (**C**).

**A,** No difference in toluidine blue staining is observed between control and *Rac1*-KO mice.

**B,** Toluidine blue staining is stronger in *Rac1/Rac3*-DKO than in control mice.

**C,** Magnified back skin photographs of *Rac3*-KO and *Rac1/Rac3-DKO* mice. *Rac1/Rac3-DKO*back skin is more strongly stained by toluidine blue than *Rac3*-KO back skin.

**Supplementary Figure 2: Keratin5 (K5) promoter functioning cells, organs, and tissues detected by X-gal staining.**

*K5-Cre^−/−^;LacZ^−/−^* and *K5-Cre^+/−^;LacZ^+/−^* mice (**A**: P1, **B-F**: P25) were fixed by 4% PFA, and then X-gal staining was performed using whole body, organs, tissues, and sections.

**A** and **B,** Whole bodies (**A**) and sections of plantar skins (**B**) were subjected to X-gal staining. Whole skin, epidermis, and sweat glands (arrowheads) of K5-Cre^+/−^;LacZ^+/−^, but not K5-Cre^*−/−*^;LacZ^*−/−*^, mice are positive for X-gal staining.

**C,** Brains and sagittal sections including the ventricle were subjected to X-gal staining. Ependymal cells of K5-Cre^+/−^;LacZ^+/−^, but not K5-Cre^*−/−*^;LacZ^*−/−*^, mice are positive to X-gal staining.

**D,** Digestive tracts (from the esophagus to rectum) and sections of the esophagus and stomach were subjected to X-gal staining. Epithelial cells in the esophagus and stomach indicated by arrows in K5-Cre^+/−^;LacZ^+/−^ mice are positive for X-gal staining.

**E,** Respiratory tract (from the trachea to the lung) and sections of trachea were subjected to X-gal staining. Tracheal glands (arrowheads) of K5-Cre^+/−^;LacZ^+/−^, but not K5-Cre^*−/−*^;LacZ^*−/−*^, mice are positive for X-gal staining.

**F,** The heart, spleen, liver, and kidney were subjected to X-gal staining. No apparent X-gal positive signal is observed either in K5-Cre^+/−^;LacZ^+/−^ or K5-Cre^*−/−*^;LacZ^*−/−*^ mice.

**Supplementary Figure 3: Differentiation of C3H/10T1/2 mesenchymal cells to adipocytes by BMP2 + FGF21.**C3H/10T1/2 cells were treated with DMI plus no factor, BMP2, FGF21, or BMP2 + FGF21. Fixed cells were stained using Oil Red O solution. Oil Red O staining was strongest in the BMP2 + FGF21 treated well (scale bar: 5 mm). Lower panels are magnified images of the upper panels. Arrows indicate the Oil Red O positive cells (scale bars: 250 μm).

**Supplementary Figure 4: Phosphorylation of Smad1/5 in intradermal white adipocyte tissue.** Intradermal white adipocyte tissue (dWAT) was obtained from *Rac3*-KO (control) and *Rac1/Rac3*-DKO mice. Lysates were subjected to SDS-PAGE followed by immunoblotting using anti-phospho-Smad1/5 and Smad1 antibodies. Phosphorylation of Smad1/5 was reduced in dWAT of *Rac1/Rac3*-DKO mice compared with that of *Rac3*-KO mice. Comparable loading of proteins was confirmed using an anti-Smad1 antibody.

